# *De Novo* Design of Peptide Binders to Conformationally Diverse Targets with Contrastive Language Modeling

**DOI:** 10.1101/2023.06.26.546591

**Authors:** Suhaas Bhat, Kalyan Palepu, Lauren Hong, Joey Mao, Tianzheng Ye, Rema Iyer, Lin Zhao, Tianlai Chen, Sophia Vincoff, Rio Watson, Tian Wang, Divya Srijay, Venkata Srikar Kavirayuni, Kseniia Kholina, Shrey Goel, Pranay Vure, Aniruddha J. Desphande, Scott H. Soderling, Matthew P. DeLisa, Pranam Chatterjee

## Abstract

Designing binders to target undruggable proteins presents a formidable challenge in drug discovery, requiring innovative approaches to overcome the lack of putative binding sites. Recently, generative models have been trained to design binding proteins via three-dimensional structures of target proteins, but as a result, struggle to design binders to disordered or conformationally unstable targets. In this work, we provide a generalizable algorithmic framework to design short, target-binding linear peptides, requiring only the amino acid sequence of the target protein. To do this, we propose a process to generate naturalistic peptide candidates through Gaussian perturbation of the peptidic latent space of the ESM-2 protein language model, and subsequently screen these novel linear sequences for target-selective interaction activity via a CLIP-based contrastive learning architecture. By integrating these generative and discriminative steps, we create a **Pep**tide **Pr**ioritization via **CLIP** (**PepPrCLIP**) pipeline and validate highly-ranked, target-specific peptides experimentally, both as inhibitory peptides and as fusions to E3 ubiquitin ligase domains, demonstrating functionally potent binding and degradation of conformationally diverse protein targets *in vitro*. Overall, our design strategy provides a modular toolkit for designing short binding linear peptides to any target protein without the reliance on stable and ordered tertiary structure, enabling generation of programmable modulators to undruggable and disordered proteins such as transcription factors and fusion oncoproteins.

## Introduction

Over 80% of pathogenic proteins are considered “undruggable” by standard small molecule inhibitors, as they often lack putative stable binding pockets.^1,2^ Targeted protein degradation (TPD) has emerged as a promising alternative to treat disease, but largely relies on small molecule warheads to bind and recruit endogenous E3 ubiquitin ligases to targets, thus still requiring the presence of accessible surface binding sites.^3^ Protein-based modalities, either as standalone binders or recruiters of effector domains, thus represent attractive platforms to target these proteins, as they do not require surface pockets for binding. As a result, numerous approaches have been developed to leverage protein-based binding domains for the design of intracellular protein degraders. These approaches, such as bioPROTACs, AdPROMs, and ubiquibodies, have relied on the existence of “off-the-shelf” binders, thus precluding degrader design to novel target substrates.^4–10^ Still, the design of new protein binders is challenging: one would either need to conduct laborious experimental screening via strategies such as yeast or phage display, or deploy structure-based computational design methods.^11–14^ Though progress is being made,^15^ these methods, such as the state-of-the-art RFDiffusion generative model, rely on three-dimensional structural information of target proteins for binder design, making design challenging to unstructured and intrinsically-disordered proteins, which make up a large fraction of the undruggable proteome.^13,14,16,17^

Recently, protein language models (pLMs) have been pre-trained on millions of natural protein sequences to generate latent embeddings that grasp relevant physicochemical, functional, and most notably, tertiary structural information.^18–21^ Even more interestingly, early results suggest that sequence-based protein transformers can produce novel protein sequences with functional capability.^22,23^ Motivated by these findings, here, we develop a pipeline to design *de novo* target-binding peptide motifs, only requiring the amino acid sequence of the target protein. Inspired by the groundbreaking work of OpenAI to generate photorealistic images from natural language descriptions via contrastive language-image pretraining (CLIP),^24^ we first leverage the ESM-2 pLM to generate distinct, but naturalistic peptide candidates by applying Gaussian noise to the embeddings of bona-fide peptide binder sequences.^18^ We then screen these candidates for target-selective interaction activity through a CLIP-based contrastive latent space, trained to jointly encode cognate peptide-protein pairs. Our resulting **Pep**tide **Pr**ioritization via **CLIP (PepPrCLIP)** pipeline integrates our generative-discriminative framework to prioritize fully novel peptide candidates with binding capacity to an input target sequence. After demonstrating competitive performance to RFDiffusion on structured targets *in silico*, we extensively validate that PepPrCLIP-prioritized “guide” peptides, when experimentally used as inhibitory peptides or fused to E3 ubiquitin ligase domains, exhibit robust, superior target engagement and modulation of diverse targets, including an enzyme, transcription factor, and undruggable fusion oncoprotein *in vitro*. Our integrative methodology thus provides a modular, CRISPR-like system for programmable TPD applications.

## Results

### A CLIP discriminator accurately captures peptide-protein binding

Recently, OpenAI released CLIP, a model that is trained to associate images and their textual descriptions by learning to rank the correct matching pairs higher than incorrect ones via contrastive learning.^24,25^ DALL-E 2, a denoising diffusion model, was then trained using the CLIP latent space to generate photorealistic images conditioned on an associated text prompt.^25^ We thus hypothesized that just as CLIP connects images to their corresponding captions using jointly-trained image and caption encoders, we can leverage a CLIP-based architecture to map target proteins to their corresponding binding peptides using jointly-trained peptide and target protein encoders. Specifically, we task our model to predict the cosine similarity between all n^2^ target and peptide pairs in a matrix, and subsequently calculate the average of the cross-entropy losses on the rows and columns of this matrix during training. We further employ Meta AI’s state-of-the-art ESM-2 model weights to generate feature-rich embeddings for cognate targets and peptides as inputs to the model (Figure 1A).^18^

**Figure 1.**
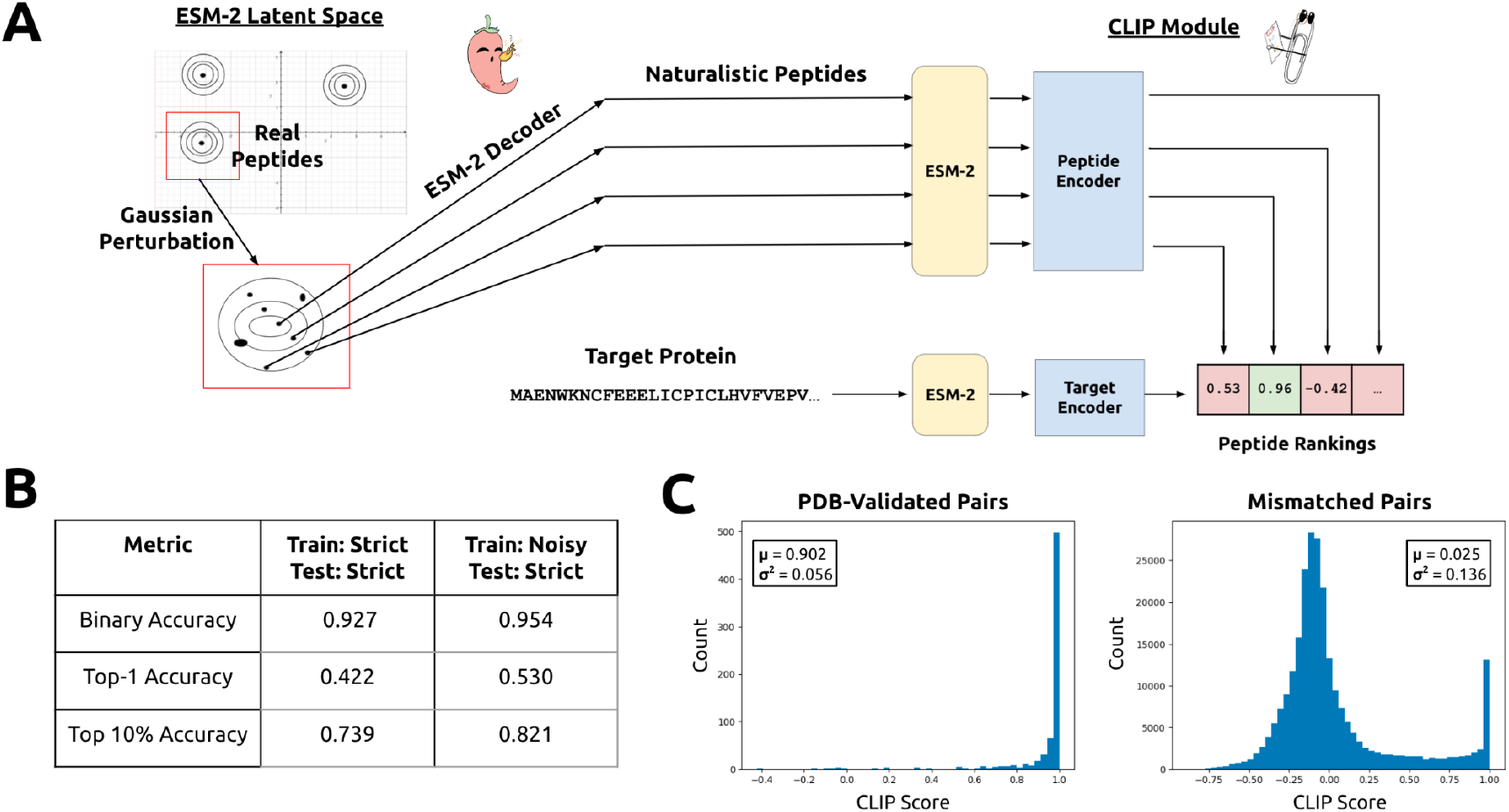
PepPrCLIP model training and evaluation. (A) Gaussian distributions centered around the ESM-2 embeddings of naturally-occurring peptides are sampled and then decoded back to amino acid sequences. A trained CLIP module, jointly encoding cognate peptide-protein embeddings, screens thousands of these peptides for specific binding activity to the target. (B) Evaluation metrics for final CLIP module. Binary accuracy is the accuracy of the model in predicting the correct binding pairs when given 2 protein-peptide pairs. Top-1 accuracy is the likelihood that for a given protein, the model has chosen the correct binding peptide, and Top 10% accuracy is the likelihood that this peptide is in the top 10% of peptides when ranked for CLIP score to this protein. Metric values are presented for the “strict” held-out test set described in the Results and Methods sections. (C) Known peptide-target sequence pairs, from the PDB-validated dataset, were scored via the trained CLIP module. Mismatched pairs, each containing a peptide and a non-matching protein, both from the validation dataset, were also scored via the CLIP module. +1 indicates a definitive binding pair while -1 indicates that the peptide and target sequences do not bind (see Methods). Scores are represented as a histogram. The mean (μ) and variance (σ^2^) are provided for each distribution.

In order to construct the dataset for training this model, we first considered every interaction structure in the RCSB PDB, and specifically filtered to create appropriate training and testing datasets. To train on a wide distribution of peptide-protein pairs, we generate a “noisy” dataset, in which we consider both strong and weaker interactions (co-crystal buried surface area >= 50 Å^2^), and consider interactions between all short proteins (< 50 AA) and another larger protein. For stringent evaluation of our model, we create a “strict” peptide-protein test dataset, in which we only include strong interactions (co-crystals with a buried surface area >= 400 Å^2^), with peptides of length <= 25 AA, and target proteins of length >= 30 AA. These parameters are equivalent to the datasets for the recent PepNN model, a deep attention model which accurately predicts peptide binding sites on target proteins.^26^ Overall, the datasets are clustered at 30% sequence identity, using MMSeqs2, ensuring significant sequence diversity between the train and test sets.^27^ After filtration and clustering, the noisy training set consists of 11,597 peptide-protein pairs, with a validation set and test set of 1241 and 1376 pairs respectively. The strict training set consists of 7388 peptide-protein pairs, with a validation set and test set of 737 and 1002 pairs respectively.

For evaluation, we use three key metrics: binary accuracy, Top-1 accuracy, and Top 10% accuracy. Binary accuracy is the accuracy of the model in predicting the correct binding pairs when given 2 protein-peptide pairs. Top-1 accuracy is the likelihood that for a given protein, the model has chosen the correct binding peptide from a batch of 64 randomly sampled peptides from the dataset, and Top 10% accuracy is the likelihood that this peptide is in the top 10% of peptides when ranked by CLIP score to this protein. Thus, Top 10% accuracy allows us to measure the model’s ability to sort a set of peptide candidates accurately. If limited resources are available for screening, a high Top 10% accuracy provides confidence that the model will have prioritized high-quality peptide candidates.

Our results demonstrate that training the CLIP model on the noisy dataset — a more difficult binding prediction task — allows for robust performance on the stricter held-out test set, exhibiting a 95.4% binary accuracy, as well as a Top 10% accuracy of 0.82 and Top-1 accuracy of 0.53. In contrast, training and testing on the strict dataset alone yields a lower 92.7% binary accuracy and poorer Top 10% and Top-1 accuracies of 0.74 and 0.42, respectively, justifying our transfer learning approach (Figure 1B). We further conducted an ablation study on the input embeddings to the CLIP model, and demonstrate that when following the two training paradigms, deployment of ESM-2 embeddings for both the peptide and target encoders is critical for high performance, as compared to a simple learned embedding or BLOSUM62 embeddings (Supplementary Figure 1),^28,29^ highlighting the importance of leveraging expressive, pre-trained pLM embeddings.

Finally, as an additional method to validate the optimized model’s ability to discriminate between peptide-protein pairs that bind and do not bind, we compare predicted CLIP scores on the ∼1000 held-out true interacting pairs from our test dataset and the ∼500,000 mismatched pairs which are assumed not to interact. As a visualization, the distribution of CLIP scores for interacting peptide-protein pairs is heavily skewed towards +1 (mean = 0.902, variance = 0.056), while mismatched non-interacting pairs have CLIP scores distributed around 0 (mean = 0.025, variance = 0.136) (Figure 1C). As a note, we observe a set of false positives in the distribution of the mismatched pairs, an expected phenomenon as natural peptides are rarely highly-specific to their target alone. Nonetheless, our results demonstrate that the CLIP model robustly selects and highly ranks target-binding peptides for a provided target sequence, thus motivating its utilization for *de novo* peptide prioritization.

### Gaussian perturbation generates naturalistic peptides for CLIP prioritization

There exist approximately 10^26^ possible 20-mer peptides — far too large of a space to exhaustively screen either *in silico* or *in vitro* for a given target. Furthermore, a small fraction of these possible peptides are likely to be biologically viable. Here, we propose a method to generate biologically naturalistic peptides. We begin with an ensemble of bona-fide, interacting peptides, which appear in co-crystals in the RCSB PDB (these peptides are taken from our training set). Next, we embed the sequences of these peptides using ESM-2. ESM-2’s latent space contains an understanding of naturalistic protein sequences — similar position in latent space corresponds to biological similarity. Thus, we expect that if we sample from the latent space in the region surrounding naturally occurring peptides, we will recover peptide sequences that are similar to those found in nature. To do this, we add noise to the embeddings of each residue of a source peptide. Noise is sampled from a standard Gaussian distribution and multiplied by a scaling factor *k* to create a perturbation, which we then add to the embedding for its respective residue. By analyzing the dependence of Hamming distance between generated peptides and their source peptides, we select values of *k* in the range 5 to 22 to maintain naturalistic peptide sequences (Figure 2A). By varying the amount of noise and the number of real peptides sampled, we generate an arbitrarily large set of biologically viable *de novo* peptides. Finally, we convert the perturbed embeddings to protein sequences using the final layer of ESM-2, acquiring new peptide sequences for CLIP screening, thus completing the end-to-end PepPrCLIP pipeline (Figure 1A). The PepPrCLIP algorithm is formally defined via pseudocode in Supplementary Figure 2.

**Figure 2.**
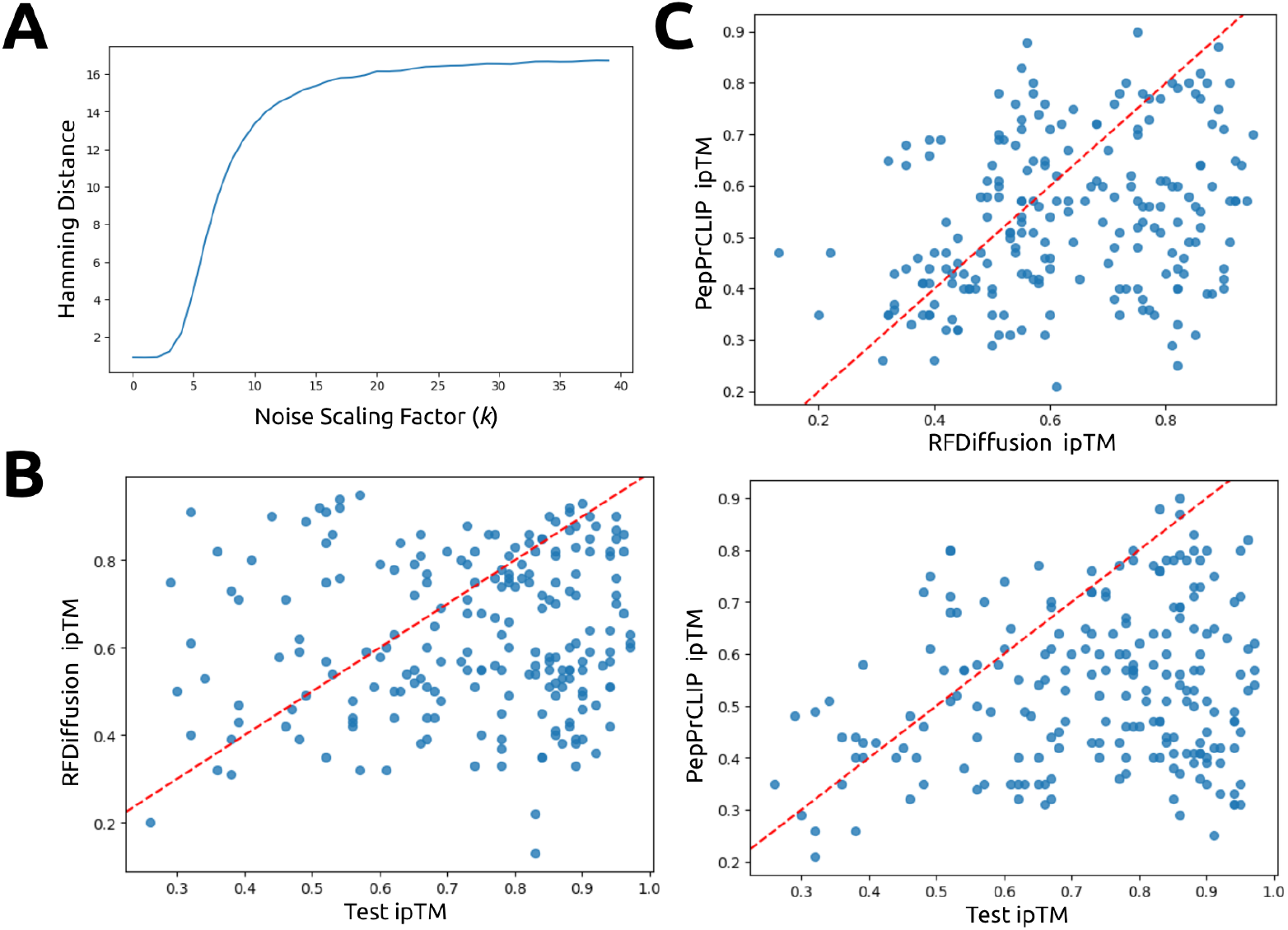
PepPrCLIP generation and *in silico* benchmarking. (A) Analysis of Hamming distance of generated peptides to source peptides vs. variance scaling factor (*k*) of embedding perturbation. We computed the mean Hamming distance between generated peptides and their source peptides as we varied the value of k. We did this for an ensemble of 100 source 18-mer peptides, creating 500 generated peptides for each. The plot of Hamming distance as we varied k is depicted in Figure 2A. We observed that at k = 5, we see mutations on the order of 1 amino acid, and mean Hamming distance increases until k = 22, at which point almost the entire sequence is mutated. Thus, we chose that k would range between 5 and 22, sampling the full range of sequence transformations. (B) *In silico* hit-rate assessment of PepPrCLIP vs. RFDiffusion. Utilizing AlphaFold-Multimer, ipTM scores were computed for both the generated and test peptides in conjunction with the target protein sequence. The entries are organized in accordance with the ipTM scores attributed to the test set peptides. The hit-rate is characterized by the generated peptides exhibiting ipTM scores ≥ those of the ground truth test peptides. (C) Post co-folding with input structured target, comparison of ipTM scores of PepPrCLIP-generated peptides to RFDiffusion-generated peptides. All ipTM scores, targets, and peptide sequences are provided in the Supplementary Data file.

### PepPrCLIP performs comparably to RFDiffusion on structured targets *in silico*

Prior to experimental testing, we computationally benchmarked the full PepPrCLIP pipeline via AlphaFold-Multimer, a massively improved method for peptide-protein docking as compared to previous docking strategies.^30,31^ Specifically, the ipTM score is a verified metric within AlphaFold2-Multimer that proxies as a potential interface binding affinity of peptide-protein complex, thus serving as a quantitative assessment of generation quality.^30,31^ To define a rigorous “hit-rate” for PepPrCLIP, we first co-folded 209 PDB-validated peptides in PepPrCLIP’s unseen, non-homologous, held-out “strict” test set and retrieved their ipTM values. We then co-folded the top PepPrCLIP-generated peptide and the top RFDiffusion-generated peptide of equivalent length with the corresponding target protein sequence (Supplementary Data).

To establish a very strict but normalized threshold on binding capacity, we calculated the hit-rate as the percentage of generated peptides that possess an ipTM score ≥ that of the corresponding test set peptide over all tested target proteins. To note, this is a very high standard, as this indicates the rate at which peptides outperform experimentally validated, high-affinity, crystallizable peptide binders. As all target proteins are PDB-validated structures likely in its training set, RFDiffusion performs strongly as expected, with an *in silico* hit-rate of 30.1% (Figure 2B). Without explicitly leveraging any structural information nor seeing these target proteins in its training set, PepPrCLIP achieves an *in silico* hit-rate of 17.7% (Figure 2B). We further calculated the percentage of peptides from each model that achieves an ipTM ≥ 0.7, demonstrating that 22% PepPrCLIP-generated peptides form stable interactions, compared to 40% of RFDiffusion-generated peptides, which has a comparative advantage on these targets.^31,32^ We also compare the ipTMs of PepPrCLIP-generated peptides and RFDiffusion-generated peptides, and find that PepPrCLIP outperforms RFDiffusion on 33% of this set of targets (Figure 2C). Overall, these results motivated us to experimentally explore PepPrCLIP’s generation quality in application-relevant settings on difficult, more conformationally diverse target proteins.

### PepPrCLIP-generated peptides inhibit the catalytic activity of UltraID

Proximity-dependent biotinylation identification (BioID) proteomics is a powerful and popular approach to study protein-protein interactions and intracellular organization. It relies on the enzymatic activity of biotin protein ligase to biotinylate proteins in proximity to a bait of interest. UltraID, a conformationally-stable enzyme (pLDDT = 96.2) evolved from the original *E. coli* BirA biotin protein ligase (Figure 3A), stands out as the smallest efficient enzyme currently available for BioID.^33^ Its high enzymatic kinetics and low background activity make it highly promising for diverse proteome discovery applications. However, employing UltraID to explore specific protein-protein interactions, either within or between cells, encounters a significant limitation: the pressing need to precisely regulate its enzymatic activity in both spatial and temporal dimensions. To overcome this obstacle, one strategy is the development of a “caged” version of UltraID, wherein it is held in an inactive state that can be released. We reasoned it may be feasible to cage UltraID by linking it with an inhibitory peptide (IP) capable of binding to the catalytic core and inhibiting its activity, until release (Figure 3B). It is noteworthy that UltraID utilizes both ATP and biotin as substrates for its biotinylating function.^33^ Therefore, the success of inhibition by the IP likely hinges on its competitive binding between IP and ATP/biotin at the active site, or by rendering the binding pocket inaccessible due to induced conformational changes.

**Figure 3.**
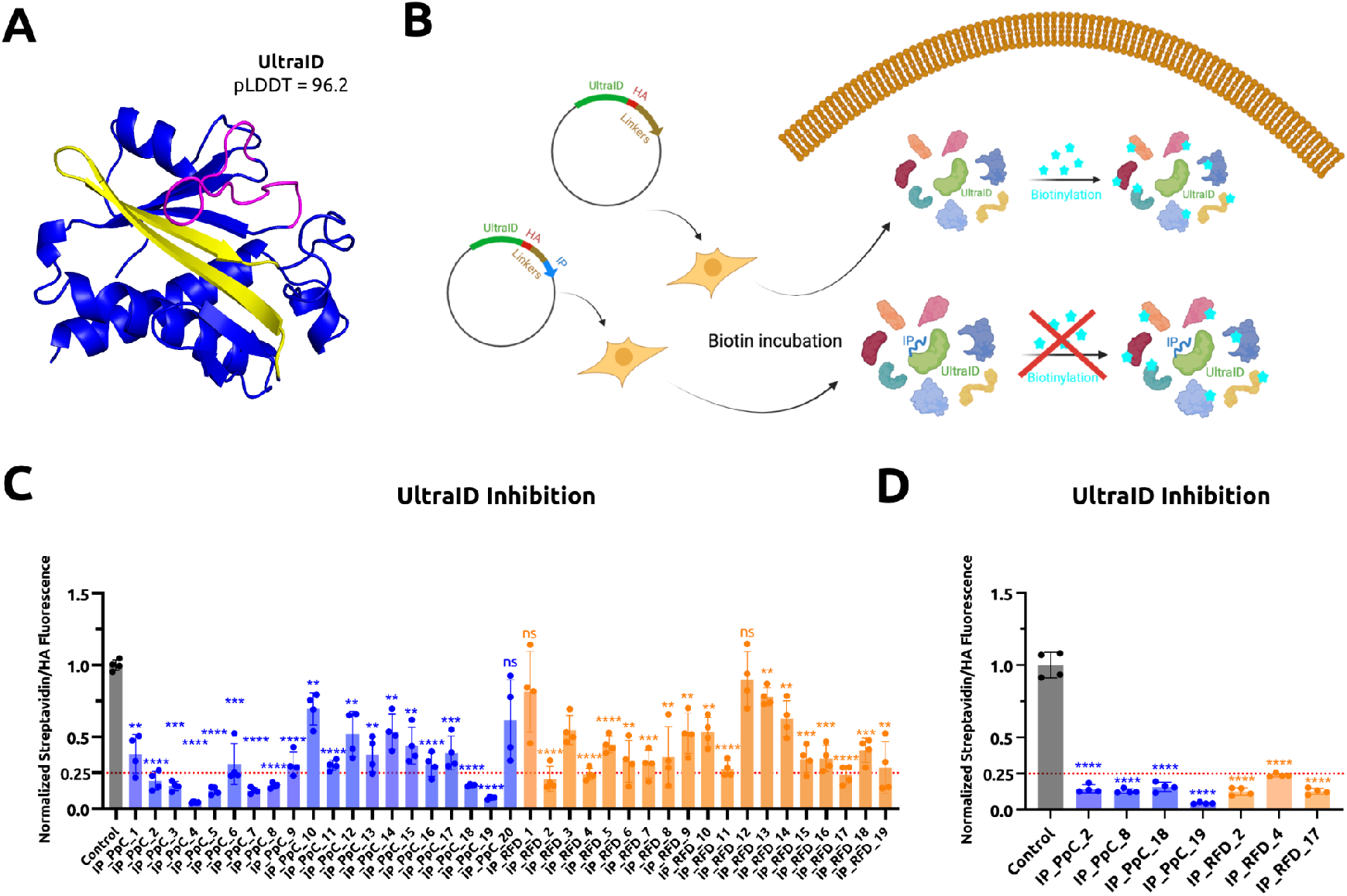
Characterization of PepPrCLIP/RFDiffusion-generated peptides for UltraID inhibition. (A) AlphaFold2-predicted structure of UltraID. The magenta region refers to hotspot amino acids 35-51 and the yellow region refers to hotspot amino acids 102-124. (B) Schematic of IP-induced inhibition of UltraID catalytic activity. Briefly, each of the PepPrCLIP/RFDiffusion-generated peptides was fused to the C-terminus of UltraID via flexible linkers and the HA epitope in a pCAG vector. The resulting plasmids were separately transfected into HEK293T cells. At 48 hours post-transfection, the cells were treated with 50 µM biotin for 30 mins before fixation and immunostaining. The control plasmid is the vector expressing UltraID but not any IP. (C) Initial screening and quantification of UltraID inhibition efficiency of PepPrCLIP (blue) and RFDiffusion (orange)-generated IPs. The inhibition efficiency was evaluated by the ratio between biotinylation level (streptavidin fluorescence level) and transfection efficiency (HA fluorescence level), which was further normalized to the control samples. Each plasmid was transfected into two wells of HEK293T cells in 24-well plates, and two fluorescence images were captured for each well. Statistical significance was determined by an unpaired two-tailed Student’s *t* test. Calculated *p* values are represented as follows: *, *p* < 0.05; **, *p* < 0.01; ***, *p* < 0.001; ****, *p* < 0.0001; ns, not significant. (D) Quantification of UltraID inhibition efficiency of 7 top candidate IPs in the independent biological replicates (*n* = 3). The experiment and statistical analysis were performed similarly as described in 3B.

Here we evaluated the ability PepPrCLIP and RFDiffusion to generate IPs targeting sequences of the UltraID catalytic pocket, which was predicted based on AlphaFold2 structural models and the known structure of BirA (Figure 3A). This was followed by experimental validation of their inhibition efficiency in cells. Providing only the target sequence of UltraID, which is not in the PDB and thus not in either PepPrCLIP’s or RFDiffusion’s training set, and confirming their catalytic pocket-targeting via AlphaFold-Multimer, we generated 20 candidate IPs (IP1-20) with PepPrCLIP (Supplementary Table 1 and Supplementary Data). Providing the AlphaFold2-predicted structure of UltraID as input and specifying the catalytic pocket as a binding hotspot, we generated 19 IPs (IP21-39) from RFDiffusion for comparison (Supplementary Table 1 and Supplementary Data). We generated plasmids expressing each of these IPs, fused to the UltraID DNA coding sequence via several flexible linkers and a hemagglutinin antibody (HA) epitope. These plasmids were then transfected into HEK293T cells, which were treated with 50 µM biotin for 30 mins before fixation and immunostaining. To assess inhibition efficiency, we evaluated the ratio between biotinylation level (streptavidin fluorescence level) and transfection efficiency (HA fluorescence level). Our results reveal that among the 19 RFDiffusion-generated IPs, 17 of them led to UltraID inhibition, and 3 (IP22, IP24, & IP37) inhibited the enzymatic activity of UltraID by more than 75 percent, highlighting its strong binder generation capability on structured targets (Figure 3B). Of the 20 PepPrCLIP-generated IPs, 19 showed successful UltraID inhibition, with 4 (IP2, IP8, IP18, & IP19) exhibiting over 75 percent inhibition efficiency (Figure 3B). Notably, the top 3 effective IPs (IP8, IP18, & IP19) were all generated by PepPrCLIP, underscoring its strong potential in designing peptides that not only bind to but also inhibit the activity of the target protein (Figure 3C). These results underlie the potential utility of PepPrCLIP in generating peptides capable of modulating the function of enzymes through predicted binding to catalytic sites.

### PepPrCLIP-generated ubiquibodies robustly bind and functionally degrade β-catenin

Based on the seminal work of Portnoff *et al*.,^8^ we recently reprogrammed the specificity of a modular human E3 ubiquitin ligase called CHIP (carboxyl-terminus of Hsc70-interacting protein) by replacing its natural substrate-binding domain, TPR, with designer “guide” peptide motifs to generate minimal, genetically-encodable ubiquibody (uAb) architectures for target protein degradation.^34^ In multiple studies, we have demonstrated the suitability of the uAb architecture to incorporate linear guide peptides designed via both structure- and sequence-based algorithms.^35–37^

To demonstrate the capacity of PepPrCLIP to design binders to targets beyond conformationally-stable, structured target proteins, we evaluated the interaction and degradation capacities of PepPrCLIP-derived peptides in the context of uAbs. For the first target, we chose β-catenin (Figure 4A), a bifunctional protein with critical roles in both cell adhesion and transcriptional regulation of the Wnt pathway whose stabilization leads to aberrant Wnt/β-catenin signaling and promotes a transcriptional program composed of many oncogenes such as *c-Myc* and *cyclin D-1*. After cloning six high-scoring PepPrCLIP-generated peptides (Supplementary Table 2) into plasmids upstream of the CHIPΔTPR catalytic domain, we transfected these plasmids into DLD1 colon cancer cells, which express wild-type β-catenin at abnormally high levels.^38^ Utilizing TOPFlash, a luciferase reporter for reliable readout of Wnt/β-catenin signaling,^39^ as an initial screening tool, we observed a statistically significant reduction in β-catenin transcriptional activity in cells transfected with β-cat_PpC_1, β-cat_PpC_2, β-cat_PpC_3, and β-cat_PpC_4 constructs compared to cells transfected with an empty vector control (Figure 4B).

**Figure 4.**
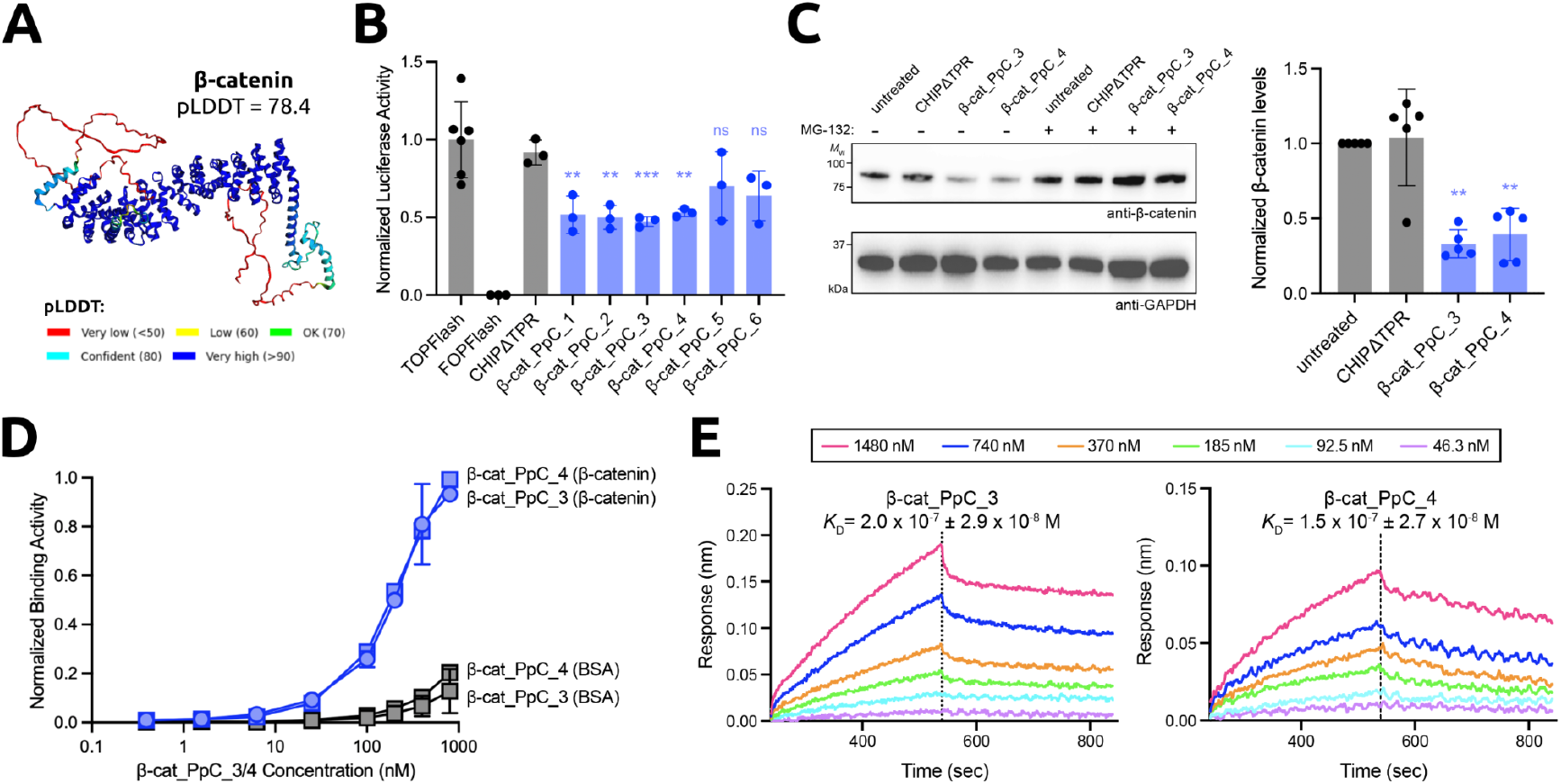
Characterization of PepPrCLIP-generated peptide-guided uAbs for β-catenin. (A) AlphaFold2-predicted structure of β-catenin. (B) TOPFlash luciferase reporter assay of Wnt/β-catenin transcriptional activity. FOPFlash reporter served as a negative control. (C) **Left:** Degradation of endogenous β-catenin in cytosolic fractions of DLD1 cells analyzed via immunoblotting with anti-β-catenin and anti-GAPDH antibodies. **Right:** Densitometry analysis of immunoblots (*n* = 3) using ImageJ software was performed to quantify β-catenin levels. Statistical significance was determined by an unpaired two-tailed Student’s *t* test. Calculated *p* values are represented as follows: **, *p* < 0.01; ***, *p* < 0.001; ns, not significant. Raw, uncropped blots are shown in Supplementary Figure 4. (D) β-catenin binding activity determined by ELISA with immobilized β-catenin. Binding to bovine serum albumin (BSA) served as a negative control. (E) Biolayer interferometry (BLI) analysis of β-cat-PpC3-CHIPΔTPR and β-cat-PpC4-CHIPΔTPR. Details on methodologies can be found in the Methods section.

To determine if this functional data was the result of endogenous β-catenin depletion, we conducted immunoblot analysis on the cytosolic fractions obtained from DLD1 cells transfected with the strong-performing β-cat_PpC_3 and β-cat_PpC_4 constructs (Figure 4C). The immunoblots revealed that these two peptide-guided uAbs promoted robust β-catenin degradation (>50%) relative to non-transfected DLD1 control cells. Importantly, degradation was proteasome-dependent, as treatment with the potent proteasome inhibitor, MG-132, blocked β-catenin degradation. We also confirmed that β-cat_PpC_3 and β-cat_PpC_4 facilitated target degradation through specific, peptide-mediated binding to β-catenin, as evidenced by ELISA (Figure 4D). Specifically, β-cat_PpC_3 and β-cat_PpC_4 exhibited strong binding activity to immobilized β-catenin with virtually no binding to the immobilized bovine serum albumin (BSA) control. Finally, to quantify binding affinity of the two peptide-guided uAbs, we performed biolayer interferometry (BLI) and determined that β-cat_PpC_3 and β-cat_PpC_4 both exhibited sub-µM binding affinity to β-catenin (*K*_D_ = 200 and 150 nM, respectively; Figure 4E), thus demonstrating the potent binding capability of PepPrCLIP-designed peptides.

### PepPrCLIP-generated uAbs robustly degrade the highly-disordered fusion oncoprotein SS18-SSX1

Synovial sarcoma belongs to a subcategory of sarcomas called soft-tissue sarcomas which accounts for 5% to 10% of all soft-tissue tumors^40^ and is more prevalent in adolescents and young adults.^41^ This disease is characterized by a singular, highly disordered oncogenic fusion protein SS18-SSX1 (Figure 5A) formed by the translocation of (X;18)(p11.2;q11.2), which leads to the fusion of the SS18 gene to one of three SSX genes (SSX1, SSX2 and rarely SSX4). In two thirds of cases SS18 is fused to the SSX1 gene.^42^ The SS18-SSX1 fusion protein is believed to function as an aberrant transcriptional regulator hijacking the activity of the normal BAF complex and driving oncogenesis.^43^ As the complete role of the fusion is not yet fully understood, targeted protein degradation of SS18-SSX1 is not only an attractive therapeutic opportunity but also a vital reagent to understand its role in the oncogenesis of synovial sarcoma.

**Figure 5.**
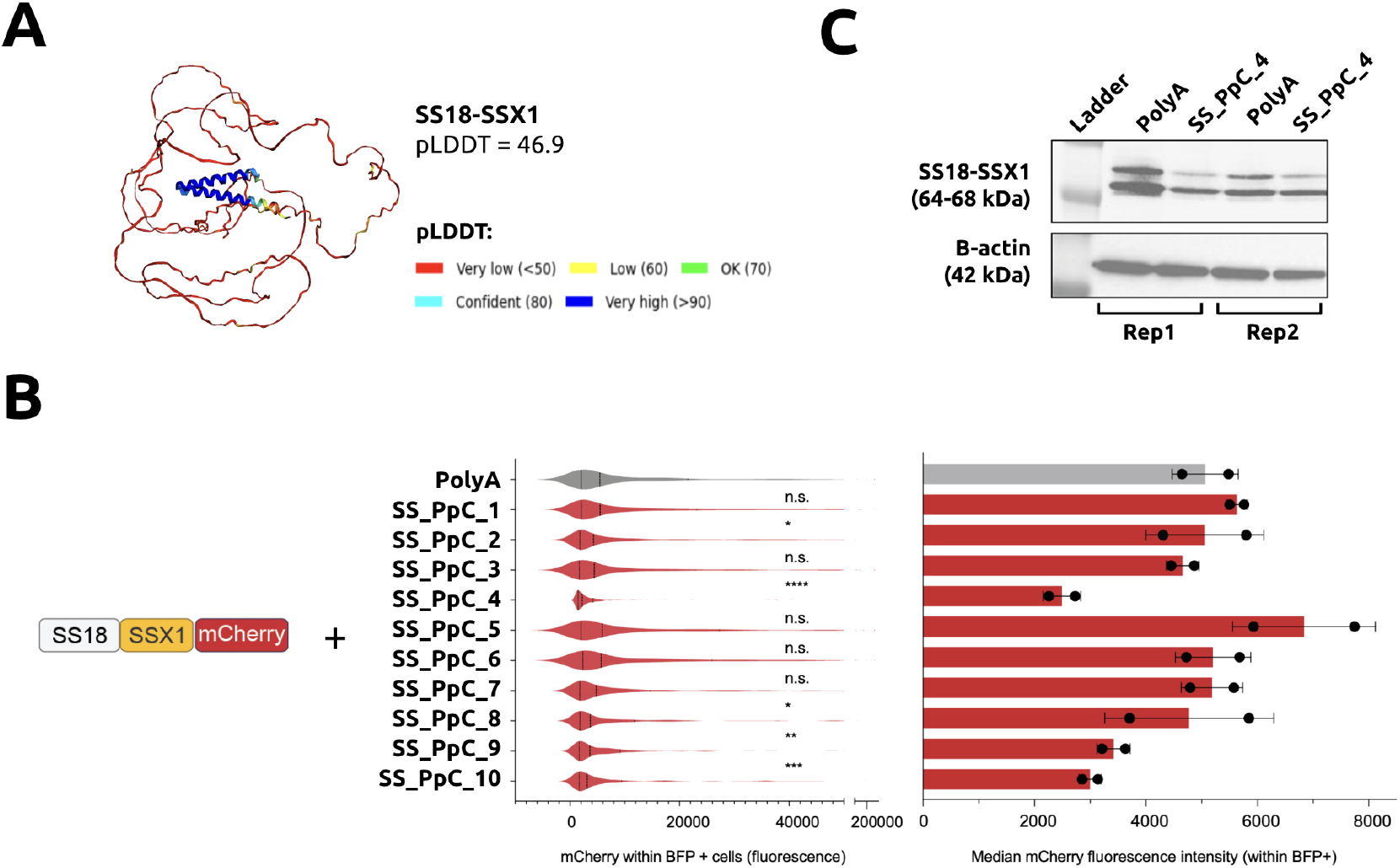
Characterization of PepPrCLIP-generated peptide-guided uAbs for SS18-SSX1 fusion. (A) AlphaFold2-predicted structure of SS18-SSX1. (B) **Left:** Schematic representation of the STY-SSX1 fusion protein fused to mCherry is shown. **Middle:** Fluorescence scale values of mCherry for SS18-SSX1 mCherry-fusion expressing cells within the peptide-expressing BFP positive fraction are plotted as a violin plot. The polyalanine (PolyA) control (gray) or each of the 10 SYT-SSX1 peptides (red) are shown (middle graph). Statistical significance was determined by an unpaired two-tailed Student’s *t* test. Calculated *p* values are represented as follows: **, *p* < 0.01; ***, *p* < 0.001; ****, *p* < 0.0001; ns, not significant. **Right:** Bar graph of median fluorescence intensity (MFI) for the PolyA control (gray) or the 10 peptides are shown. Samples were treated in independent biological replicates (*n* = 2). Gating strategy is described in Supplementary Figure 3. (C) Immunoblot to probe endogenous SS18-SSX1 fusion in HS-SYII synovial sarcoma cells treated with SS-PpC_4 3 days post transient transfection. β-actin was used as loading control. Experiments represent independent biological replicates (*n* = 2). Raw, uncropped blots are shown in Supplementary Figure 4.

To assess whether PepPrCLIP-designed peptides can induce target degradation SS18-SSX1 protein within the uAb architecture, we first cloned the fusion in-frame with the mCherry fluorescent protein and ectopically expressed the construct in HEK293T cells. We also simultaneously co-transfected 10 individual peptides designed with PepPrCLIP (Supplementary Table 3), which were fused to CHIPΔTPR-IRES-TagBFP, and monitored the expression of the SS18-SSX1 fusion via the linked mCherry reporter (see Schematic in Figure 5B). Of the peptides predicted by PepPrCLIP to bind to SS18-SSX1, 10 were tested and of those, SS_PpC_4 significantly (*p* value <0.0001, Wilcoxon Rank Sum Test) reduced SS18-SSX1-mCherry fluorescence as monitored via flow cytometry (Figure 5B). Next, we tested the effect of SS_PpC_4 overexpression on endogenous SYT-SSX1 fusion protein levels. For this, we transfected the synovial sarcoma cell line HS-SY-II with the SS_PpC_4 peptide or the polyalanine peptide control. Strikingly, SS_PpC_4 peptide overexpression led to a strong reduction in SS18-SSX1 protein levels as assessed by immunoblotting with a fusion-specific antibody (Figure 5C). These findings corroborate the results obtained from flow cytometry and provide additional evidence of the efficacy of SS_PpC_4 in inducing targeted degradation of the fusion protein, showing that the algorithm is effective for previously intractable, disordered targets such as large fusion oncoproteins.

Overall, these results emphasize the simple, CRISPR-like programmability of PepPrCLIP-designed uAbs, which may prove to be most useful for research applications, replacing more labor intensive systems such as dTAG^44^ for studying phenotypic effects of target protein degradation.

## Discussion

Here, we describe a purely sequence-based approach for the design of *de novo* binding peptides, which we use to programmably bind, inhibit, and degrade conformationally-diverse substrate proteins *in vitro* in a CRISPR-like manner. While previous works have utilized structure-based models for binding disordered motifs,^45^ contrastive learning for predicting protein function,^46^ and language models for isolating peptidic fragments from interacting protein sequences,^35^ PepPrCLIP represents a novel example of leveraging sequence-based language models for fully *de novo* binder generation. Overall, the promising results from our approach motivate future development of methods utilizing a candidate generator and contrastive discriminator screening models for molecular design. We are also exploring alternative generative processes, such as techniques leveraging databases of antibody CDRs or human protein motifs and autoregressive models, such as GPT, Llama2, and Mamba,^47–49^ and even extending to post-translationally modified (PTM) target sequences using PTM-aware pLM embeddings.^50^ For future work, we plan to expand our *in silico* screening strategies to optimize other key properties, including selectivity and binding affinity to undruggable proteins via fine-tuning on specific target classes, such as fusion oncoproteins.^51^ In total, our two-step method represents a first protein binder design module from target sequence alone, without the requirement of 3D structural information. Our easy-to-use model, coupled with our unique experimental modalities, thus establish a more modular, CRISPR-like alternative to protein targeting, and open the door for programmable proteome editing and screening applications.

## Methods

### Dataset Generation

The datasets for this paper were generated by mining the RCSB PDB (https://www.rcsb.org/) for verified protein-protein and peptide-protein interaction structures. Two datasets were curated: 1) A “strict” peptide-protein dataset, which filters for strong interactions (co-crystals with a buried surface area >= 400 Å^2^), with peptides of length <= 25 AA, and target proteins of length >= 30 AA. 2) A “noisy” dataset which contains both strong interactions and filters the PDB-derived data for weaker interactions (co-crystal buried surface area >= 50 Å^2^) as well as interactions between all proteins (<50 AA) paired with a larger protein (>50 AA). Protein sequences were then clustered using MMseqs2 connected component clustering at 0.3 minimum sequence identity to group homologous proteins.^27^ Additional parameters used for data filtration can be found in Abdin, et al.^26^ After filtration and clustering, the noisy training set consists of 11,597 peptide-protein pairs, with a validation set and test set of 1241 and 1376 pairs respectively. The strict training set consists of 7388 peptide-protein pairs, with a validation set and test set of 737 and 1002 pairs respectively.

### CLIP Architecture and Training

The CLIP architecture was adapted from Radford, et al.^24^ Encoders were trained such that the cosine similarity between target protein embeddings and peptide embeddings, defined as

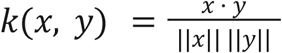

is near 1 for target-peptide pairs which do bind to each other, and is near -1 for target-peptide pairs which do not bind to each other.

Next, the target and peptide encoders were trained on batches of *n* pairs of targets and peptides which are known to interact from the datasets described above. Target and peptide sequences were encoded by their respective encoders, producing target embeddings *r*_1_,…, *r*_*n*_, and peptide embeddings *p*_1_, …, *p*_*n*_. The cosine similarity between all *n*^2^ target and peptide pairs is computed in a matrix *K*, defined such that:

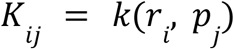

Cosine similarities are interpreted as softmax logits. We define two cross-entropy losses, one on the matrix rows and one the matrix columns:

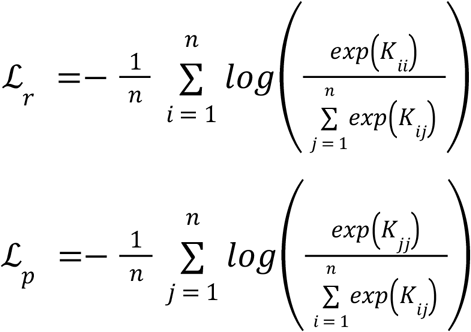

ℒ_*r*_ represents the loss on the model’s ability to predict the correct target given a single peptide, while ℒ_*p*_ represents the loss on the model’s ability to predict the correct peptide given a single target. By using these cross-entropy losses, we implicitly assumed that the *n*^*2*^ − *n* target-peptide pairs in the batch which are not known interactions do not bind at all. While this may not be a completely accurate assumption, it is approximately true.

The target encoder and peptide encoder have identical architectures. First, the input sequence is embedded through a frozen ESM-2 model, producing a *l* × *e*_*i*_ input embedding (where *l* is the input sequence length and *e*_*i*_ is the dimension of the ESM-2 embedding). Next, input embedding is averaged over the sequence length *l*, yielding an embedding vector of length *e*_*i*_ for both peptides and proteins. *h* MLP layers were applied with ReLU activation on the embedding vector to obtain the output embedding, which is a vector of length *e*_*o*_. For the final model: *e*_*i*_ = 1280, *e*_*o*_ = 320, and *h* = 2. Combining the vector embeddings of peptide and protein via dot product yields a CLIP score, valued between -1 and 1. The model is trained such that binding pairs of peptides and proteins have high CLIP scores. Thus, the model constructs a latent space of dimension *e*_*o*_ in which vector similarity corresponds to predicted binding. It can be noted that this latent space further provides a notion of protein-protein similarity, and peptide-peptide similarity, as proteins with similar binding behavior will be mapped in proximity to one another within the phase space (and the same is true for peptides). Notably, since the hidden MLP layers operate on the embedding averaged over the length dimension, the encoder has minimal dependence on the sequence length. This is particularly important for allowing the peptide encoder to generalize to a broad range of peptide lengths.

Model hyperparameters were tuned manually with the goal of optimizing validation loss. Models were trained on a single Nvidia A100 80 GB GPU via Google CoLab using an Adam optimizer for 56 epochs across the training set using early stopping (monitoring the validation loss on the “strict” dataset) with a patience of 6. All presented *in silico* metrics were done with train/validation/test splits indicated in the Results and Dataset Generation section. For experimental peptide derivation, a model trained on the entire dataset was employed to ensure maximum performance.

### Generation of peptide candidate sequences

Candidate peptides are generated using all of the peptides in the training set. Each peptide is embedded within ESM-2 pLM (https://huggingface.co/docs/transformers/model_doc/esm) in PyTorch. For a given peptide embedding, the variance (σ^2^) is computed over all dimensions of the embedding. For each residue in the source peptide, noise is sampled from the standard Normal distribution *N*(0, *k*^2^) and multiplied by the variance σ^2^ to create a perturbation *p*_*r_i*_. This perturbation is added to the embedding for its respective residue. The noise scaling factor *k* can be varied to produce peptides that are closer or farther from the source peptides in the ESM phase space. At inference time, source peptides are randomly sampled from the training set (usually on the order of 100 to 1000). For each source peptide, 1000 peptides are generated through the noising method described above, varying the value of *k* in order to sample at various distances from the source peptide. Finally, these peptides (on the order of 100,000) are fed into the CLIP model, and ranked based on predicted binding to a user-provided target sequence.

For generation of peptides for this study, *k* values in the range 5 to 22 were selected. We computed the mean Hamming distance between generated peptides and their source peptides as we varied the value of k. We did this for an ensemble of 100 source 18-mer peptides, creating 500 generated peptides for each. The plot of Hamming distance as we varied k is depicted in Figure 2A. We observed that at k = 5, we see mutations on the order of 1 amino acid, and mean Hamming distance increases until k = 22, at which point almost the entire sequence is mutated. Thus, we chose that k would range between 5 and 22, sampling the full range of sequence transformations. All peptide sequences evaluated in this study can be found in Supplementary Tables 1 and 2.

### AlphaFold-Multimer benchmarking

AlphaFold-Multimer_v3 was used in ColabFold: https://colab.research.google.com/github/sokrypton/ColabFold/blob/main/AlphaFold2.ipynb.^52^ The predicted interface TM-score (ipTM) was extracted from the post-run output files and plotted via Matplotlib. 3D images shaded with pLDDT scores were extracted from the ColabFold interface.^52^ An ipTM score above 0.7 indicates effective binding from previous studies.^31^

### RFDiffusion Generation

In parallel to the PepPrCLIP approach, RFDiffusion was employed to design peptide binders for both cases. For the given test set, RFDiffusion was tasked with generating one peptide binder per target protein, matching the length specified by the ground truth binders. The predicted structures were then converted into sequences using ProteinMPNN. For the selected critical proteins, RFDiffusion and ProteinMPNN generated candidate binders of equivalent length to test set peptides for a given target. RFDiffusion inference code on ColabFold can be found here: https://colab.research.google.com/github/sokrypton/ColabDesign/blob/v1.1.1/rf/examples/diffusion.ipynb

### Generation of plasmids

All uAb plasmids were generated from the standard pcDNA3 vector, harboring a cytomegalovirus (CMV) promoter and a C-terminal P2A-eGFP cassette. An Esp3I restriction site was introduced immediately upstream of the CHIPΔTPR CDS and flexible GSGSG linker via the KLD Enzyme Mix (NEB) following PCR amplification with mutagenic primers (Genewiz). For uAb assembly, oligos for candidate peptides were annealed and ligated via T4 DNA Ligase into the Esp3I-digested uAb backbone. Assembled constructs were transformed into 50 µL NEB Turbo Competent *Escherichia coli* cells, and plated onto LB agar supplemented with the appropriate antibiotic for subsequent sequence verification of colonies and plasmid purification (Genewiz). The SS18-SSX1 fusion gene fused directly upstream and in frame with the mCherry fluorescent protein and separated by a flexible glycine-serine linker (pLV-Puro-EF1a-SS18-SSX1-3XGGGGS-mCherry) was constructed using Vectorbuilder. In addition, 10 top-scoring peptides predicted to bind to the SS18-SSX1 protein using the PepPrCLIP were cloned in frame with the CHIPΔTPR in an altered pcDNA3.1 expression vector that co-expresses IRES-TagBFP.

### UltraID inhibition assay

HEK293T cells were seeded in 24-well plates (3×10^4^/well) with coverslips at the bottom and transfected at ∼50% confluency. 0.25 μg UltraID-IP plasmids were transfected into cells with Polyethylenimine Max (PEI Max, Polysciences 24765) in Opti-MEM (Gibco 11058021). At 3 days post transfection, the cells were treated with 50 µM biotin (Sigma-Aldrich B4501) in Dulbecco’s phosphate-buffered saline (PBS, ThermoFisher 14190144) for 30 min. Prior to immunocytochemistry, the cells were first fixed with fixation buffer (4% (v/v) paraformaldehyde (PFA, Sigma-Aldrich 158127), 4% (w/v) sucrose in 1x PBS) at 4°C for 10 min. Then the cells were gently shaken in 1x PBS twice at RT for 5 min, and then permeabilized with 0.25% (v/v) Triton X-100 (Sigma-Aldrich T9284) in 1x PBS at room temperature (RT) for 7 min. For immunocytochemistry, the cells were gently shaken in blocking buffer (5% (v/v) normal goat serum (Jackson ImmunoResearch), 0.2% (v/v) Triton X-100 in 1x PBS) at RT for 1 h, followed by overnight incubation in the blocking buffer with primary antibodies at 4°C. On the following day, the cells were gently shaken in washing buffer (0.1% (v/v) Triton X-100 in 1x PBS) thrice at RT for 5 min, and in the blocking buffer with secondary antibodies at RT for 1 h, and in the washing buffer thrice at RT for 5 min. The coverslips were transferred onto glass slides with FluroSave Reagent (Millipore). Fluorescent images were captured using an upright microscope (Zeiss Axio Imager 2) with a 20x objective lens. Image analysis was performed with ImageJ. Antibodies used and the corresponding dilutions are as follows: rat-anti-HA (1:500, Millipore 11867423001), goat-anti-rat 488 (1:500, Invitrogen A11006), and streptavidin 568 conjugate (1:500, Invitrogen S11226).

### TOPFlash assay

DLD1 cells per well were seeded on a white-bottom 96-well plate (1×10^4^/well; ThermoFisher 136101) 24 h before transfection. On the day of transfection, each well was transfected with Lipofectamine 3000 (ThermoFisher L3000001) and 100 ng of plasmid DNA in a ratio of TOPFlash (Addgene 12456)/FOPFlash (Addgene 12457): Renilla (Addgene, #87121): uAbs = 1:0.1:3. After 48 h of incubation, cells were lysed and the firefly and Renilla luminescence signals were measured sequentially by the dual-luciferase reporter kit (Promega E1910). Plates were read on a microplate reader (Tecan). The luciferase activities were measured and normalized against the control Renilla activities.

### Flow cytometry analysis

25,000 HEK293T fibroblasts per well were seeded in a 48 well plate and cultured in DMEM supplemented with 10% FBS, 2 mM L-glutamine as well as 50 u/ml penicillin/streptomycin and incubated in 5% CO_2_ at 37°C. After 24 h, cells were co-transfected with 0.2 mg of the SS18-SSX1-mCherry plasmid and 0.2 mg of each uAb encoding construct plasmid DNA in triplicates using Lipofectamine 2000 (ThermoFisher 11668019) as per manufacturer’s directions. Media was changed 24 h post-transfection and cells were analyzed on day 3 for presence of mCherry in the BFP+ gate via a flow cytometer (LSR Fortessa, BD Biosciences). The gating strategy is described in Supplementary Figure 3.

### Cell fractionation and immunoblotting

Detection of β-catenin in DLD1 cells was performed as follows. Harvested cells treated with uAbs or empty control plasmid were detached via 0.05% (w/v) trypsin-EDTA and washed twice with ice-cold 1x PBS. The cytosolic fractions were isolated from lysates using a Subcellular Protein Fractionation Kit (ThermoFisher, 78840) per the manufacturer’s instructions. Specifically, ice-cold cytosolic extraction buffer was added to the cell pellet, the mixture was placed at 4 °C for 10 min with gentle shaking followed by centrifugation at 500 × g for 10 min at 4 °C. The supernatant was collected immediately to a pre-chilled PCR tube and placed on ice followed by immunoblotting or stored at −20 °C for future use. Protein concentration was quantified using the Pierce BCA Protein Assay Kit (ThermoFisher 23225). An equivalent amount of total protein was loaded into 4–20% Mini-PROTEAN Protein Gels (BioRad 4561094) and separated by electrophoresis. Immunoblotting was performed according to standard protocols. Briefly, proteins were transferred to poly(vinylidene fluoride) (PVDF) membranes (Millipore IPVH00010), blocked with 5% (w/v) nonfat dry milk (Carnation) in 1× tris-buffered saline (TBS) with 0.05% (v/v) Tween 20 (TBST) at room temperature for 1 hour, washed three times with TBST for 5 min, and probed with rabbit anti-β-catenin antibody (Cell Signaling 8480S; diluted 1:1,000) or rabbit anti-GAPDH (Abcam ab8245; diluted 1:5,000). The blots were washed again three times with TBST for 5 min each and then probed with a secondary antibody, donkey anti-rabbit-horseradish peroxidase (HRP) (Abcam 7083; diluted 1:2,500), for 1 h at room temperature. Blots were detected by chemiluminescence using a ChemiDoc MP imager (Bio-Rad). Densitometry analysis of protein bands in immunoblots was performed using ImageJ. Intensity data for the uAb bands was normalized to band intensity for empty plasmid control. For the SYT-SSX1 experiments, HS-SYII synovial sarcoma cells were plated in a 6-well plate (250,000/well). The SS_PpC_4 uAb construct was transfected with Lipofectamine 3000 as per manufacturer’s directions. On day 3 post transfection, cells were washed with 1x PBS twice and lysed with ice-cold RIPA Lysis and Extraction Buffer (Thermo Scientific 89900) along with protease inhibitor cocktail (Thermo Scientific, Halt Protease Inhibitor Cocktail (100X) 78429). Whole cell protein was quantitated using Pierce BCA Protein Assay Kit (Thermo 23225). Protein lysates were subjected to gel electrophoresis using pre-cast Bolt 4–12% Bis-Tris Plus Gels (Invitrogen NW04127BOX) followed by nitrocellulose membrane transfer. Membranes were probed using anti-Rabbit SS18-SSX1 antibody (Cell Signaling 72364, diluted 1:1000) followed by incubation with anti-rabbit secondary conjugated with horseradish peroxidase. Protein bands were detected with the Super Signal Femto reagent (Thermo Scientific, 34095) with an anti-β-actin antibody (Cell Signaling 3700, diluted 1:1000) serving as loading control. Unprocessed, uncropped blots are shown in Supplementary Figure 4.

### Protein expression and purification

All purified uAb constructs were obtained from cultures of *E. coli* BL21(DE3) cells carrying pET28a-based plasmids. Cells were grown in Luria-Bertani (LB) medium until optical density at 600 nm reached 0.3-0.5, after which 0.1 mM isopropyl β-D-1-thiogalactopyranoside (IPTG) was added to induce protein expression. The culture was further incubated in the shaker at 16 °C for 48 h. Cells were harvested by centrifugation at 8000 rpm for 30 min at 4 °C and lysed using an EmulsiFlex-C5 high-pressure homogenizer (Avestin). Lysates were centrifuged at 15,000 rpm for 25 min at 4 °C to separate the soluble and insoluble fractions. Clarified lysates containing 6xHis-tagged proteins were subjected to gravity-flow nickel-affinity purification using HisPur Ni-NTA resin (ThermoFisher 88223) following the manufacturer’s protocols. The final purity of all proteins was confirmed by Coomassie-blue staining of SDS-PAGE gels.

### ELISA

The enzyme-linked immunosorbent assay (ELISA) was conducted as follows: 96-well plates (Corning, 9018) were coated with 2 μg/mL of β-catenin (Biomatik RPU40704) diluted in 1x PBS (pH 7.4) at a volume of 50 µL per well at 4 °C overnight. Next, the plates were washed with 1x PBST (PBS, 0.1% (v/v) Tween 20) three times and then blocked with 200 µL of blocking buffer (3% bovine serum albumin (BSA) in 1xTBST) per well overnight at 4 °C. Purified uAb constructs were biotinylated using EZ-Link Biotin (ThermoFisher 21335) following the manufacturer’s protocol. The biotinylated uAb constructs were then serially diluted in 1x PBS and added to the ELISA plates for 1 h at room temperature with gentle shaking. The plates were then washed three times with 1x PBST and incubated for 1 h at room temperature with HRP-conjugated streptavidin (ThermoFisher, Cat # N100; diluted 1:20,000), with shaking at 450 rpm. Following an additional three washes with 1x PBST, 100 µL of 3,3’-5,5’-tetramethylbenzidine substrate (1-Step Ultra TMB-ELISA; ThermoFisher, 34029) was added to each well, and the plates were incubated at room temperature in darkness. The reaction was quenched by adding 100 µL of 2 M H_2_SO4, and absorbance was quantified at a wavelength of 450 nm utilizing a FilterMax F5 microplate spectrophotometer (Agilent).

### Biolayer interferometry analysis

The dissociation constant (*K*_D_) for the purified uAbs to β-catenin was measured by biolayer interferometry (BLI) assays using an Octet RH16 instrument (Sartorius) at 30°C. Kinetic analysis was performed using streptavidin (SA) biosensor tips (Sartorius 18-5020) as follows: (1) sensor check: 30 s immersion in Octet buffer (1x PBS containing 0.2 μm filtered 0.1% BSA and 0.05% (v/v) Tween 20); (2) loading: 60 s immersion in Octet buffer supplemented with 1 μg/ml biotinylated-uAbs; (3) baseline: 60 s immersion in Octet buffer; (4) association: 300 s immersion in solution with β-catenin at concentration: 1480 nM, 740 nM, 370 nM, 185 nM, 92.5 nM, and 46.3 nM; (5) dissociation: 300 s immersion in Octet buffer. The kinetic data was analyzed using Octet Analysis Studio software v12.2.2.26 (Sartorius).

### Statistical analysis and reproducibility

All data were reported as average values with error bars representing standard deviation (SD). For individual samples performed in independent biological triplicates (n=3) or above, statistical significance was determined by unpaired *t* test (\**p* < 0.05, \*\**p* < 0.01; \*\*\**p* < 0.001; \*\*\*\**p* < 0.0001). All graphs were generated using Prism 9 for MacOS version 9.2.0. No data were excluded from the analyses. The experiments were not randomized. The investigators were not blinded to allocation during experiments and outcome assessment.

## Author Contributions

S.B. and K.P. designed the PepPrCLIP architecture, curated peptide-protein data, trained the CLIP module, and evaluated trained models. S.B. developed the peptide generation method. K.P. developed the CLIP module, with optimization done by S.B. J.M. performed UltraID experiments. L.H. and T.Y. performed β-catenin experiments and data analysis. R.I. performed SS18-SSX1 experiments. L.H., L.Z. and R.W. performed additional uAb screening experiments, with assistance from T.W. and D.S. T.C. and S.V. performed *in silico* peptide generation and AlphaFold2-Multimer benchmarking, with assistance from V.S.K., K.K., S.G., and P.V. A.J.D. supervised SS18-SSX1 experiments. S.H.S. supervised UltraID experiments. M.P.D supervised β-catenin experiments. S.B., K.P., A.J.D., S.H.S., L.H., T.C., T.Y., M.P.D. and P.C. wrote the manuscript, with input from all authors. P.C. conceived, designed, directed, and supervised the study.

## Data and Materials Availability

All data needed to evaluate the conclusions in the paper are present in the paper and supplementary tables and figures, as well as the Supplementary Data file. All code and Colab notebooks to run PepPrCLIP can be freely accessed by the academic community at: https://huggingface.co/ubiquitx/pepprclip, after signing a non-commercial, research-only academic license. All raw and processed data have been deposited to the Zenodo repository: https://zenodo.org/doi/10.5281/zenodo.10971077.

## Acknowledgements

We thank UbiquiTx, Inc. for access to compute resources and peptide-protein CLIP architectures for the manuscript. We thank Sarah Pertsemlidis, Elena Haarer, and Vivian Yudistyra for technical assistance.

## Declarations

P.C. is supported by the National Cancer Institute (Award #R21CA278468), The Hartwell Foundation, the Wallace H. Coulter Foundation, and institutional startup funds. A.J.D. is supported by NIH grants CA262746 and P30 CA030199. S.H.S. is supported by NIH grants MH111684 and MH126954. UbiquiTx, Inc. funded the CLIP model development and experiments related to β-catenin silencing, and also owns rights to PepPrCLIP.

## References

1. Behan, F. M. et al. Prioritization of cancer therapeutic targets using CRISPR–Cas9 screens. Nature 568, 511–516 (2019).

2. Dang, C. V., Reddy, E. P., Shokat, K. M. & Soucek, L. Drugging the ‘undruggable’ cancer targets. Nat. Rev. Cancer 17, 502–508 (2017).

3. Zhao, L., Zhao, J., Zhong, K., Tong, A. & Jia, D. Targeted protein degradation: mechanisms, strategies and application. Signal Transduction and Targeted Therapy 7, 1–13 (2022).

4. Lim, S. et al. bioPROTACs as versatile modulators of intracellular therapeutic targets including proliferating cell nuclear antigen (PCNA). Proc. Natl. Acad. Sci. U. S. A. 117, 5791–5800 (2020).

5. VanDyke, D., Taylor, J. D., Kaeo, K. J., Hunt, J. & Spangler, J. B. Biologics-based degraders - an expanding toolkit for targeted-protein degradation. Curr. Opin. Biotechnol. 78, 102807 (2022).

6. Bery, N., Miller, A. & Rabbitts, T. A potent KRAS macromolecule degrader specifically targeting tumours with mutant KRAS. Nat. Commun. 11, 1–14 (2020).

7. Cong, F., Zhang, J., Pao, W., Zhou, P. & Varmus, H. A protein knockdown strategy to study the function of beta-catenin in tumorigenesis. BMC Mol. Biol. 4, 10 (2003).

8. Portnoff, A. D., Stephens, E. A., Varner, J. D. & DeLisa, M. P. Ubiquibodies, synthetic E3 ubiquitin ligases endowed with unnatural substrate specificity for targeted protein silencing. J. Biol. Chem. 289, 7844–7855 (2014).

9. Stephens, E. A. et al. Engineering Single Pan-Specific Ubiquibodies for Targeted Degradation of All Forms of Endogenous ERK Protein Kinase. ACS Synth. Biol. 10, 2396–2408 (2021).

10. Simpson, L. M. et al. Inducible Degradation of Target Proteins through a Tractable Affinity-Directed Protein Missile System. Cell Chem Biol 27, 1164–1180.e5 (2020).

11. Könning, D. & Kolmar, H. Beyond antibody engineering: directed evolution of alternative binding scaffolds and enzymes using yeast surface display. Microb. Cell Fact. 17, 32 (2018).

12. Alfaleh, M. A. et al. Phage Display Derived Monoclonal Antibodies: From Bench to Bedside. Front. Immunol. 11, 1986 (2020).

13. Cao, L. et al. Design of protein-binding proteins from the target structure alone. Nature 605, 551–560 (2022).

14. Bennett, N. R. et al. Improving de novo protein binder design with deep learning. Nat. Commun. 14, 1–9 (2023).

15. Liu, C. et al. Diffusing protein binders to intrinsically disordered proteins. bioRxiv (2024) doi:10.1101/2024.07.16.603789.

16. Watson, J. L. et al. Broadly applicable and accurate protein design by integrating structure prediction networks and diffusion generative models. bioRxiv (2022) doi:10.1101/2022.12.09.519842.

17. Gainza, P. et al. De novo design of protein interactions with learned surface fingerprints. Nature 617, 176–184 (2023).

18. Lin, Z. et al. Evolutionary-scale prediction of atomic-level protein structure with a language model. Science 379, 1123–1130 (2023).

19. Elnaggar, A. et al. ProtTrans: Toward Understanding the Language of Life Through Self-Supervised Learning. IEEE Trans. Pattern Anal. Mach. Intell. 44, 7112–7127 (2022).

20. Elnaggar, A. et al. Ankh ☥: Optimized protein language model unlocks general-purpose modelling. bioRxiv (2023) doi:10.1101/2023.01.16.524265.

21. Brandes, N., Ofer, D., Peleg, Y., Rappoport, N. & Linial, M. ProteinBERT: a universal deep-learning model of protein sequence and function. Bioinformatics 38, 2102–2110 (2022).

22. Madani, A. et al. Large language models generate functional protein sequences across diverse families. Nat. Biotechnol. 1–8 (2023).

23. Ferruz, N., Schmidt, S. & Höcker, B. ProtGPT2 is a deep unsupervised language model for protein design. Nat. Commun. 13, 1–10 (2022).

24. Radford, A. et al. Learning transferable visual models from natural language supervision. (2021) doi:10.48550/ARXIV.2103.00020.

25. Ramesh, A., Dhariwal, P., Nichol, A., Chu, C. & Chen, M. Hierarchical text-conditional image generation with CLIP latents. (2022) doi:10.48550/ARXIV.2204.06125.

26. Abdin, O., Nim, S., Wen, H. & Kim, P. M. PepNN: a deep attention model for the identification of peptide binding sites. Communications Biology 5, 1–10 (2022).

27. Steinegger, M. & Söding, J. MMseqs2 enables sensitive protein sequence searching for the analysis of massive data sets. Nat. Biotechnol. 35, 1026–1028 (2017).

28. Henikoff, S. & Henikoff, J. G. Amino acid substitution matrices from protein blocks. Proc. Natl. Acad. Sci. U. S. A. 89, 10915–10919 (1992).

29. ElAbd, H. et al. Amino acid encoding for deep learning applications. BMC Bioinformatics 21, 235 (2020).

30. Evans, R. et al. Protein complex prediction with AlphaFold-Multimer. bioRxiv (2021) doi:10.1101/2021.10.04.463034.

31. Johansson-Åkhe, I. & Wallner, B. Improving peptide-protein docking with AlphaFold-Multimer using forced sampling. Front Bioinform 2, 959160 (2022).

32. Tsaban, T. et al. Harnessing protein folding neural networks for peptide–protein docking. Nat. Commun. 13, 1–12 (2022).

33. Kubitz, L. et al. Engineering of ultraID, a compact and hyperactive enzyme for proximity-dependent biotinylation in living cells. Communications Biology 5, 1–14 (2022).

34. Chatterjee, P. et al. Targeted intracellular degradation of SARS-CoV-2 via computationally optimized peptide fusions. Communications Biology 3, 1–8 (2020).

35. Brixi, G. et al. SaLT&PepPr is an interface-predicting language model for designing peptide-guided protein degraders. Communications Biology 6, 1–10 (2023).

36. Palepu, K. et al. Design of Peptide-Based Protein Degraders via Contrastive Deep Learning. bioRxiv (2022) doi:10.1101/2022.05.23.493169.

37. Chen, T. et al. PepMLM: Target sequence-conditioned generation of peptide binders via masked language modeling. (2023) doi:10.48550/ARXIV.2310.03842.

38. Nusse, R. & Clevers, H. Wnt/β-Catenin Signaling, Disease, and Emerging Therapeutic Modalities. Cell 169, 985–999 (2017).

39. Korinek, V. et al. Constitutive transcriptional activation by a beta-catenin-Tcf complex in APC-/-colon carcinoma. Science 275, 1784–1787 (1997).

40. Mastrangelo, G. et al. Incidence of soft tissue sarcoma and beyond: a population-based prospective study in 3 European regions. Cancer 118, 5339–5348 (2012).

41. Sbaraglia, M., Bellan, E. & Dei Tos, A. P. The 2020 WHO Classification of Soft Tissue Tumours: news and perspectives. Pathologica 113, 70–84 (2021).

42. Hale, R., Sandakly, S., Shipley, J. & Walters, Z. Epigenetic Targets in Synovial Sarcoma: A Mini-Review. Front. Oncol. 9, 1078 (2019).

43. McBride, M. J. et al. The SS18-SSX Fusion Oncoprotein Hijacks BAF Complex Targeting and Function to Drive Synovial Sarcoma. Cancer Cell 33, 1128–1141.e7 (2018).

44. Nabet, B. et al. The dTAG system for immediate and target-specific protein degradation. Nat. Chem. Biol. 14, 431–441 (2018).

45. Wu, K. et al. De novo design of modular peptide-binding proteins by superhelical matching. Nature 616, 581–589 (2023).

46. Yu, T. et al. Enzyme function prediction using contrastive learning. Science 379, 1358–1363 (2023).

47. Brown, T. B. et al. Language Models are Few-Shot Learners. (2020) doi:10.48550/ARXIV.2005.14165.

48. Touvron, H. et al. Llama 2: Open foundation and fine-tuned chat models. (2023) doi:10.48550/ARXIV.2307.09288.

49. Gu, A. & Dao, T. Mamba: Linear-time sequence modeling with selective state spaces. (2023) doi:10.48550/ARXIV.2312.00752.

50. Peng, Z., Schussheim, B. & Chatterjee, P. PTM-Mamba: A PTM-Aware Protein Language Model with Bidirectional Gated Mamba Blocks. bioRxiv (2024) doi:10.1101/2024.02.28.581983.

51. Vincoff, S. et al. FusOn-pLM: A Fusion Oncoprotein-Specific Language Model via Focused Probabilistic Masking. bioRxiv (2024) doi:10.1101/2024.06.03.597245.

52. Mirdita, M. et al. ColabFold: making protein folding accessible to all. Nat. Methods 19, 679–682 (2022).

