## Supplementary Information for "*De Novo* Design of Peptide Binders to Conformationally Diverse Targets with Contrastive Language Modeling"

**Supplementary Figures**

1. Ablation study of CLIP input embeddings.
2. Pseudocode of PepPrCLIP algorithm.
3. Example gating strategy for flow cytometry of SS18-SSX1 degradation analysis for Figure 5B.
4. Raw, uncropped Western blots for Figure 4C and 5C.

**Supplementary Tables**

1. Peptide sequences for UltraID testing.
2. Peptide sequences for β-catenin testing.
3. Peptide sequences for SS18-SSX1 degradation experiments.

**A**

**Train: Strict, Test: Strict**

| **Peptide Embedding** | ESM-2 | ESM-2 | nn.Embedding | nn.Embedding | BLOSUM62 | BLOSUM62 |
|---|---|---|---|---|---|---|
| **Protein Embedding** | ESM-2 | nn.Embedding | ESM-2 | nn.Embedding | ESM-2 | BLOSUM62 |
| **Binary Accuracy** | 0.927 | 0.629 | 0.617 | 0.653 | 0.493 | 0.548 |
| **Top-1 Accuracy** | 0.422 | 0.040 | 0.046 | 0.039 | 0.023 | 0.016 |
| **Top 10% Accuracy** | 0.739 | 0.196 | 0.200 | 0.198 | 0.114 | 0.114 |

**B**

**Train: Noisy, Test: Strict**

| **Peptide Embedding** | ESM-2 | ESM-2 | nn.Embedding | nn.Embedding | BLOSUM62 | BLOSUM62 |
|---|---|---|---|---|---|---|
| **Protein Embedding** | ESM-2 | nn.Embedding | ESM-2 | nn.Embedding | ESM-2 | BLOSUM62 |
| **Binary Accuracy** | 0.954 | 0.514 | 0.510 | 0.634 | 0.541 | 0.594 |
| **Top-1 Accuracy** | 0.530 | 0.010 | 0.016 | 0.043 | 0.013 | 0.0355 |
| **Top 10% Accuracy** | 0.821 | 0.087 | 0.095 | 0.178 | 0.087 | 0.122 |

**Supplementary Figure 1. Ablation study of CLIP input embeddings.** Evaluation metrics for the CLIP module provided the indicated input embeddings with the specified datasets for training and testing. Input peptides and proteins were either embedded with ESM-2 or with the PyTorch class nn.Embedding, which uses a simple lookup table to maps an index value to a weight matrix of a certain dimension: (https://pytorch.org/docs/stable/generated/torch.nn.Embedding.html). Binary accuracy is the accuracy of the model in predicting the correct binding pairs when given 2 protein-peptide pairs. Top-1 accuracy is the likelihood that for a given protein, the model has chosen the correct binding peptide, and Top 10% accuracy is the likelihood that this peptide is in the top 10% of peptides when ranked for binding affinity to this protein. Metric values are presented for the “strict” held-out test set described in the Results and Methods sections.

**A**

**Algorithm 1** CLIP Training Loop

**Input:** Batch of $n$ randomly pairs of receptors and peptides ($receptor\_msa$, $peptide$)
**Output:** Trained receptor and peptide encoders
**for** $i = 1$ to $n$ **do**
  $\boldsymbol{r_i} \leftarrow$ **ReceptorEncoder**($receptor_i$)
  $\boldsymbol{p_i} \leftarrow$ **PeptideEncoder**($peptide_i$)
**end for**
Initialize matrix $\boldsymbol{K} \in \mathbb{R}^{n \times n}$
**for** $i = 1$ to $n$ **do**
  **for** $j = 1$ to $n$ **do**
    $K_{ij} \leftarrow k(\boldsymbol{r_i}, \boldsymbol{p_j}) = \frac{\boldsymbol{r_i} \cdot \boldsymbol{p_j}}{\|\boldsymbol{r_i}\| \|\boldsymbol{p_j}\|}$
  **end for**
**end for**
$\mathcal{L}_r \leftarrow -\frac{1}{n} \sum_{i=1}^{n} \log \left( \frac{\exp(K_{ii})}{\sum_{j=1}^{n} \exp(K_{ij})} \right)$
$\mathcal{L}_p \leftarrow -\frac{1}{n} \sum_{j=1}^{n} \log \left( \frac{\exp(K_{jj})}{\sum_{i=1}^{n} \exp(K_{ij})} \right)$
$\mathcal{L} \leftarrow \frac{1}{2}(\mathcal{L}_r + \mathcal{L}_p)$
**Optimize** step on Receptor and Peptide Encoders using $\mathcal{L}$

**B**

**Algorithm 2** Generate and Score Peptide Sequences

$\boldsymbol{r} \leftarrow$ **ReceptorEncoder**($receptor$)
$peptide\_scores \leftarrow \{\}$
$sampled\_peptides \leftarrow$ randomly sample $num\_base\_peps$ from $base\_peptides$
**for** each $peptide \in sampled\_peptides$ **do**
  embedding $\leftarrow$ **ESM_Embed**($peptide$)
  **for** k $\in$ $variances$ **do**
    **for** $j$ from 1 to $num\_samples\_per\_base$ **do**
      $z \leftarrow$ Sample a matrix $Z \in \mathbb{R}^{m \times n}$
      where each $Z_{ij} \sim N(0, k^2)$ for all $i, j$
      noised_embedding $\leftarrow$ embedding $+ z \cdot$ **Var**($embedding$)
      $new\_peptide \leftarrow$ **ESM_Final_Layer**(embedding)
      $\boldsymbol{p} \leftarrow$ **PeptideEncoder**($new\_peptide$)
      $new\_peptide\_score \leftarrow k(\boldsymbol{r}, \boldsymbol{p}) = \frac{\boldsymbol{r} \cdot \boldsymbol{p}}{\|\boldsymbol{r}\| \|\boldsymbol{p}\|}$
      Append ($new\_peptide$, $new\_peptide\_score$) to $generated\_peptides$
    **end for**
  **end for**
**end for**
**return** $generated\_peptides$

**Supplementary Figure 2. Pseudocode of PepPrCLIP algorithm.** (A) CLIP training loop. Peptide and target encoders are jointly trained on ESM-2 embeddings to predict high cosine similarities between known peptide-receptor embedding pairs and low cosine similarities for all other pairs. (B) Generation and scoring of peptide sequences. A large sample of training-set peptides are embedded in the ESM-2 latent space. For each "source peptide" in the training set, many peptides (e.g. 1000) are created by sampling from a standard Gaussian, and varying the noise parameter *k*. These peptides are then fed into CLIP for scoring. Critically, the generation process is entirely unconditioned on the target, or any information about the target's binding partners.

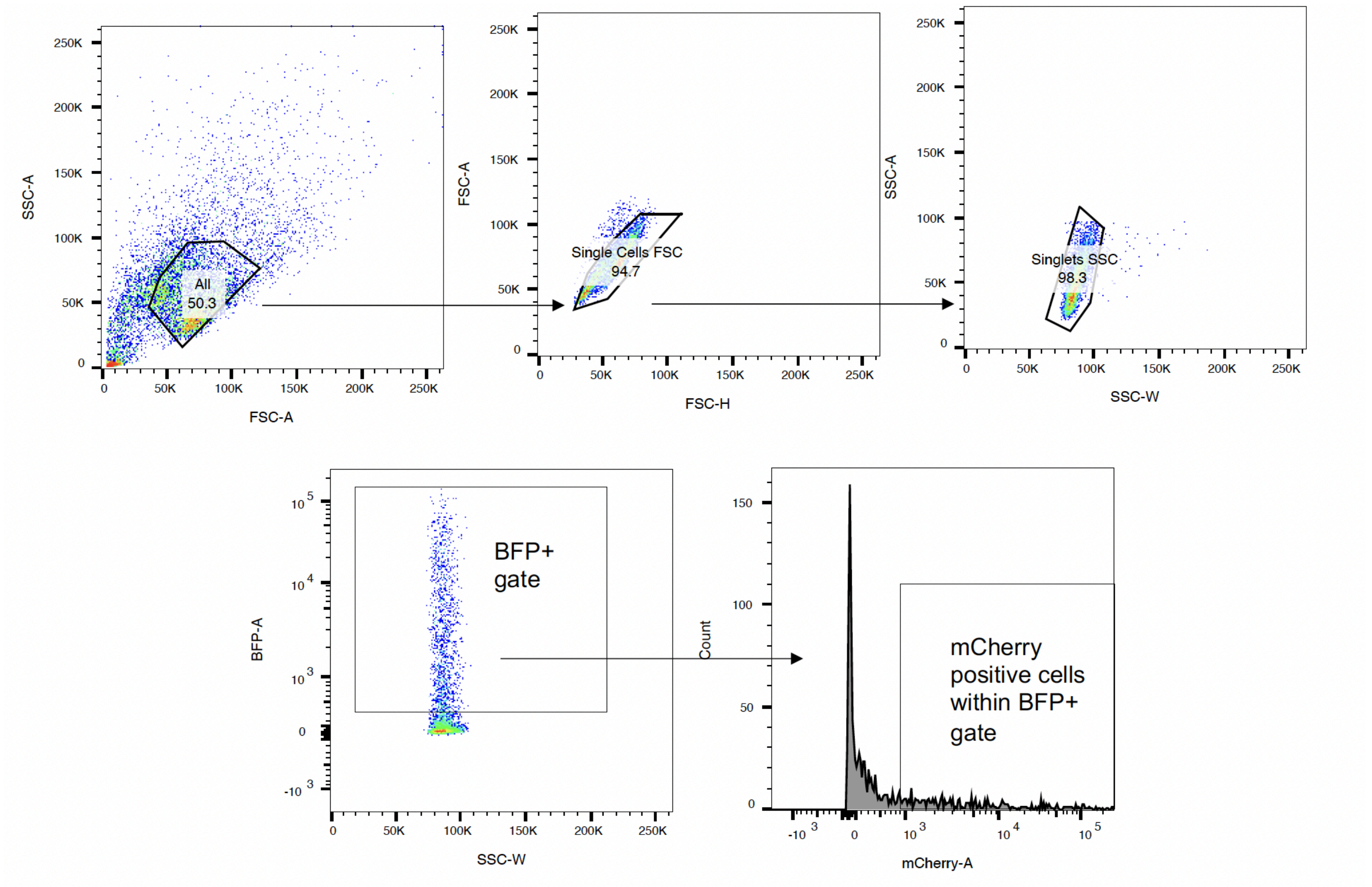


**Supplementary Figure 3. Example gating strategy for flow cytometry of SS18-SSX1 degradation analysis for Figure 5B.**

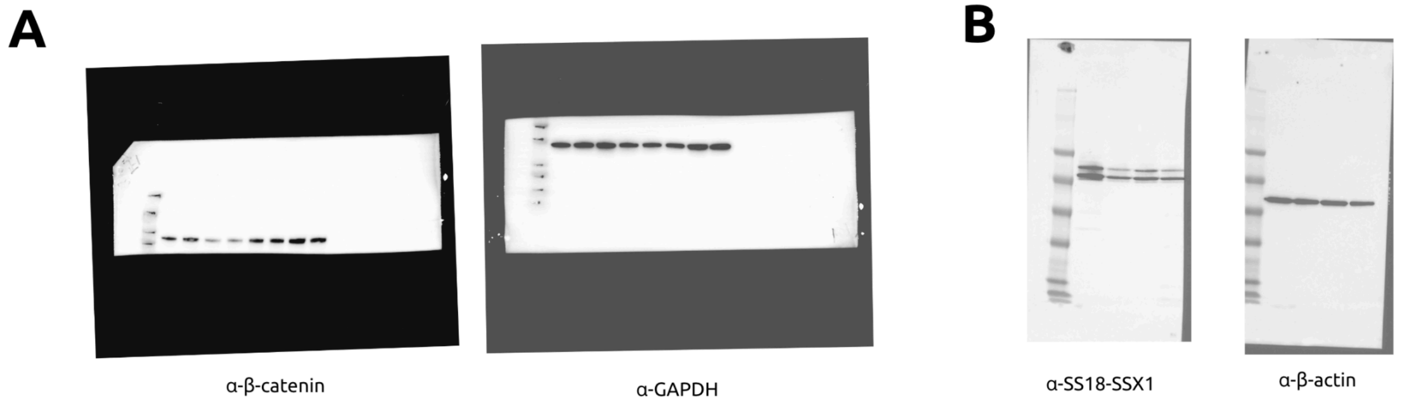


**Supplementary Figure 4. Raw, uncropped Western blots for Figure 4C and 5C.** A) Figure 4C Western blots DLD1 cytosolic fraction lysate treated with indicated primary antibodies. B) Figure 5C Western blots of HS-SY-II lysate treated with indicated primary antibodies. Western blotting details can be found in the Methods section.

**Supplementary Table 1. Peptide sequences for UltraID testing.**

**UltraID Sequence:**
MDFKNLIWLKEVDSTQERLKEWNVSYGTALVADRQTKGRGGPGRKWLSQEGGLYFSFLLNPKEFENLLQLPVLGLSVSEALEEITEIPFSLKWPNDVYFQEKKVSGVLCELSKDKLIVGIGINVNQREIPEEIKDRATTLYEITGKDWDRKEVLLKVLKRISENLKKFKEK

| uAb Name | Peptide Sequence | Derivation Method | Peptide Score (CLIP Score for PpC/ RMSD for RFD) |
|---|---|---|---|
| IP1 | REGGTQRELMDRPEWYK | PepPrCLIP | 0.72572327 |
| IP2 | VSCDGMPQFDMKFPWLKQG | PepPrCLIP | 0.70803845 |
| IP3 | DGGQTNDKGIPVNMKPMQQ | PepPrCLIP | 0.70665252 |
| IP4 | QIIKDLIEAMGFDKFKWKGE | PepPrCLIP | 0.70595598 |
| IP5 | IMTKRKTTVHEKKSLMPS | PepPrCLIP | 0.70294487 |
| IP6 | KIGWNKHQPFPVDFLKPI | PepPrCLIP | 0.76725882 |
| IP7 | WNQSKLSKQGNENNYNHKTK | PepPrCLIP | 0.75147784 |
| IP8 | DNNHCSWLYGYWEWTLWGED | PepPrCLIP | 0.73040438 |
| IP9 | LKKQTMMFEVLPMKLRWD | PepPrCLIP | 0.72898626 |
| IP10 | NEPDINMENSLNLNPNP | PepPrCLIP | 0.72232258 |
| IP11 | YKTLHQKSLWYSIEGIK | PepPrCLIP | 0.71636546 |
| IP12 | YEDMRETLDPWVTNQGIAP | PepPrCLIP | 0.71523684 |
| IP13 | YGDWKDPSDLEGDFEENF | PepPrCLIP | 0.71494853 |
| IP14 | SNPWLKPTWDDPECPYQK | PepPrCLIP | 0.71054471 |
| IP15 | VGTHTPDTNGWKIGLDH | PepPrCLIP | 0.70977223 |
| IP16 | WWAIKDFKDCIYTKSGKSE | PepPrCLIP | 0.70622885 |
| IP17 | RVWSMHEEKDRYPLKRCF | PepPrCLIP | 0.70541561 |
| IP18 | MVPKVIGLGPCYVTSMLIP | PepPrCLIP | 0.70321536 |
| IP19 | EMIDENRRRVLVKVVLFP | PepPrCLIP | 0.70292401 |
| IP20 | PVCPEKPPKLQLQQAEG | PepPrCLIP | 0.7009151 |
| IP21 | KVRVFQKENPDGTVETEVVVE | RFDiffusion | 6.1875453 |
| IP22 | AAAALERLLQQLAAALAAGEL | RFDiffusion | 13.7183361 |
| IP23 | ETKWLYQEQEP | RFDiffusion | 14.4984789 |
| IP24 | KLLETRLQTGEKPRLLALVEG | RFDiffusion | 17.0711517 |
| IP25 | VPLVNQEGKEL | RFDiffusion | 20.7622719 |

| | | | |
|---|---|---|---|
| IP26 | TKVNIVTQTAPSGKKLSLTLL | RFDiffusion | 22.2421799 |
| IP27 | KKLRLIEERDGKVIEKLVVIG | RFDiffusion | 23.4596691 |
| IP28 | KKKDWLGVEYRDGKTTVKYYT | RFDiffusion | 23.957489 |
| IP29 | KRVRETDEETGEVREVEVEVP | RFDiffusion | 24.1955204 |
| IP30 | ETVWEIEESLAASGAPAKELR | RFDiffusion | 28.8947525 |
| IP31 | KKLITLVFDL | RFDiffusion | 0.75644505 |
| IP32 | GLKLETIVVE | RFDiffusion | 0.75838602 |
| IP33 | LVLEFDAEEVKERLEKE | RFDiffusion | 0.97315383 |
| IP34 | KITKTIVLDL | RFDiffusion | 1.10380983 |
| IP35 | MKLYFEFYEK | RFDiffusion | 1.19073594 |
| IP36 | EKKKTTVLVEVIDVKVP | RFDiffusion | 1.92825592 |
| IP37 | KKVVYFEFYE | RFDiffusion | 1.98646462 |
| IP38 | EKEKKLEEILKKLKELA | RFDiffusion | 5.95443916 |
| IP39 | KKLRVEVKYEELEDILS | RFDiffusion | 9.69598675 |

**Supplementary Table 2. Peptide sequences for β-catenin testing.** Only PepPrCLIP-generated peptides were experimentally tested, as RFDiffusion-generated peptides demonstrate poor RMSD scores.

**β-catenin Sequence:**
MATQADLMELDMAMEPDRKAAVSHWQQQSYLDSGIHSGATTTAPSLSGKGNPEEEDVDTSQVLYEWEQGFS
QSFTQEQVADIDGQYAMTRAQRVRAAMFPETLDEGMQIPSTQFDAAHPTNVQRLAEPSQMLKHAVVNLINYQ
DDAELATRAIPELTKLLNDEDQVVVNKAAVMVHQLSKKEASRHAIMRSPQMVSAIVRTMQNTNDVETARCTAGT
LHNLSHHREGLLAIFKSGGIPALVKMLGSPVDSVLFYAITTLHNLLLHQEGAKMAVRLAGGLQKMVALLNKTNVK
FLAITTDCLQILAYGNQESKLIILASGGPQALVNIMRTYTYEKLLWTTSRVLKVLSVCSSNKPAIVEAGGMQALGL
HLTDPSQRLVQNCLWTLRNLSDAATKQEGMEGLLGTLVQLLGSDDINVVTCAAGILSNLTCNNYKNKMMVCQV
GGIEALVRTVLRAGDREDITEPAICALRHLTSRHQEAEMAQNAVRLHYGLPVVVKLLHPPSHWPLIKATVGLIRNL
ALCPANHAPLREQGAIPRLVQLLVRAHQDTQRRTSMGGTQQQFVEGVRMEEIVEGCTGALHILARDVHNRIVIR
GLNTIPLFVQLLYSPIENIQRVAAGVLCELAQDKEAAEAIEAEGATAPLTELLHSRNEGVATYAAAVLFRMSEDKP
QDYKKRLSVELTSSLFRTEPMAWNETADLGLDIGAQGEPLGYRQDDPSYRSFHSGGYGQDALGMDPMMEHE
MGGHHPGADYPVDGLPDLGHAQDLMDGLPPGDSNQLAWFDTDL

| uAb Name | Peptide Sequence | Derivation Method | Peptide Score (CLIP Score for PpC/ RMSD for RFD) |
|---|---|---|---|
| βcat_PpC_1 | KEPDDLMIPAFLRKYAN | PepPrCLIP | 0.995235324 |
| βcat_PpC_2 | KEDDYPDIPAFLRKQAD | PepPrCLIP | 0.990422189 |
| βcat_PpC_3 | EESDDLYIPAFLRKQLD | PepPrCLIP | 0.989606261 |
| βcat_PpC_4 | DFIPPYVDFPPRLPIFR | PepPrCLIP | 0.985456049 |
| βcat_PpC_5 | YDDDFDIKLLRKRPSKPL | PepPrCLIP | 0.983283103 |
| βcat_PpC_6 | MRRLDPKAWFEYDVDLSL | PepPrCLIP | 0.979800761 |
| βcat_RFD_1 | QEEEARRVVSQVQSAAA | RFDiffusion | 28.8681583 |
| βcat_RFD_2 | QADLMELIMERRKAAVK | RFDiffusion | 20.4560814 |
| βcat_RFD_3 | LSMLKMEDLKAAISQAQ | RFDiffusion | 29.7054539 |
| βcat_RFD_4 | MAALMEAREADRKAVLS | RFDiffusion | 30.0191288 |
| βcat_RFD_5 | DLDKILEKTEIQSKVQAQ | RFDiffusion | 31.5413609 |
| βcat_RFD_6 | DLAMAMEREALLEVQQQQ | RFDiffusion | 29.3066273 |

**Supplementary Table 3. Peptide sequences designed for SS18-SSX1.** Only PepPrCLIP-generated peptides were experimentally tested, as the target pLDDT was < 50.

**SS18-SSX1 Sequence:**
MSVAFAAPRQRGKGEITPAAIQKMLDDNNHLIQCIMDSQNKGKTSECSQYQQMLHTNLVYLATIADSNQNMQS
LLPAPPTQNMPMGPGGMNQSGPPPPPRSHNMPSDGMVGGGPPAPHMQNQMNGQMPGPNHMPMQGPGP
NQLNMTNSSMNMPSSSHGSMGGYNHSVPSSQSMPVQNQMTMSQGQPMGNYGPRPNMSMQPNQGPMMH
QQPPSQQYNMPQGGGQHYQGQQPPMGMMGQVNQGNHMMGQRQIPPYRPPQQGPPQQYSGQEDYYGDQ
YSHGGQGPPEGMNQQYYPDGHNDYGYQQPSYPEQGYDRPYEDSSQHYYEGGNSQYGQQQDAYQGPPPQ
QGYPPQQQQYPGQQGYPGQQQGYGPSQGGPGPQYPNYPQGQGQQYGGYRPTQPGPPQPPQQRPYGYD
QIMPKKPAEDENDSKGVSEASGPQNDGKQLHPPGKANISEKINKRSGPKRGKHAWTHRLRERKQLVIYEEISD
PEEDDE

| Sample Name | Peptide Sequence | Derivation Method | Peptide Score (CLIP Score for PpC/ RMSD for RFD) |
|---|---|---|---|
| PolyA | MAAAAAAAAAAAAAAA | N/A | N/A |
| SS_PpC_1 | YPREDWHHWHQKCPKR | PepPrCLIP | 0.983730375 |
| SS_PpC_2 | MKDGQHWHFRKHLYLGYH | PepPrCLIP | 0.964908599 |
| SS_PpC_3 | VPPPPPLPDGPPWPNL | PepPrCLIP | 0.958866834 |
| SS_PpC_4 | LWWGMAKYCAAKLTCMC | PepPrCLIP | 0.956480264 |
| SS_PpC_5 | AGPVAGHVEPPPPWPPGA | PepPrCLIP | 0.947725594 |
| SS_PpC_6 | MICHRNCHLARDVFKR | PepPrCLIP | 0.947692453 |
| SS_PpC_7 | MDVTITVNGKVVAKP | PepPrCLIP | 0.941476821 |
| SS_PpC_8 | YCHPTRHEHLIRPRRTDP | PepPrCLIP | 0.941453755 |
| SS_PpC_9 | MPPKLCWCNVERHRYYH | PepPrCLIP | 0.940890729 |
| SS_PpC_10 | DHALRPQRHWCHLNH | PepPrCLIP | 0.937198162 |
| SS_RFD_1 | QLLFELIRAALAAYLA | RFDiffusion | 12.8079729 |
| SS_RFD_2 | SVEELQQQLAEIKAKLAA | RFDiffusion | 8.43772411 |
| SS_RFD_3 | SLLELLKLQVEAVSAA | RFDiffusion | 6.24589157 |
| SS_RFD_4 | LSSLLAVLLQMLLLQLS | RFDiffusion | 9.16260147 |
| SS_RFD_5 | SLEELLRQLEEVKARLLA | RFDiffusion | 2.83086419 |
| SS_RFD_6 | SLLEILSSILSTLLAA | RFDiffusion | 3.84221196 |
| SS_RFD_7 | SLLEQLLAEIVAAAQ | RFDiffusion | 11.2154684 |
| SS_RFD_8 | SLLKEILEQIKAFVEAYK | RFDiffusion | 5.03273439 |
| SS_RFD_9 | STIEELKEQLETLRATF | RFDiffusion | 2.46420264 |
| SS_RFD_10 | SAAIEAALKRLLEAL | RFDiffusion | 3.53702259 |
